# PACS-1 and Adaptor Protein-1 Mediate ACTH Trafficking to the Regulated Secretory Pathway

**DOI:** 10.1101/446229

**Authors:** Brennan S. Dirk, Christopher End, Emily N. Pawlak, Logan R. Van Nynatten, Rajesh Abraham Jacob, Bryan Heit, Jimmy D. Dikeakos

## Abstract

The regulated secretory pathway is a specialized form of protein secretion found in endocrine and neuroendocrine cell types. Pro-opiomelanocortin (POMC) is a pro-hormone that utilizes this pathway to be trafficked to dense core secretory granules (DCSGs). Within this organelle, POMC is processed to multiple bioactive hormones that play key roles in cellular physiology. However, the complete set of cellular membrane trafficking proteins that mediate the correct sorting of POMC to DCSGs remain unknown. Here, we report the roles of the phosphofurin acidic cluster sorting protein – 1 (PACS-1) and the clathrin adaptor protein 1 (AP-1) in the targeting of POMC to DCSGs. Upon knockdown of PACS-1 and AP-1, POMC is readily secreted into the extracellular milieu and fails to be targeted to DCSGs.

## INTRODUCTION

In addition to the constitutive secretion of proteins into the extracellular milieu, endocrine and neuroendocrine cells also possess a specialized form of secretion termed “regulated secretion” [1-3]. Interestingly, regulated secretion is often preceded by the specific targeting of unprocessed pro-hormones to a specialized subcellular storage organelle termed the dense core secretory granule (DCSG) [4, 5]. Within DCSGs, pro-hormones and their substrates can be processed into their active forms. Upon receiving the appropriate physiological stimulus, vesicles containing DCSG resident cargo fuse with the plasma membrane, and are secreted into the extracellular milieu [5].

Pro-opiomelanocortin (POMC) is a pro-hormone that undergoes proteolytic cleavage to produce smaller active peptide hormones within DCSGs [6-8]. This activity requires the co-targeting of both POMC, and pro-hormone convertase (PC) enzymes to DCSGs where they mediate the conversion of POMC to its active forms ([9] and reviewed in [10]). The products generated from POMC cleavage by PCs include peptide hormones, such as α/β/γ melanotropin; which regulate skin pigmentation [11], β-endorphin; an endogenous opioid [12], and adrenocorticotropic hormone (ACTH); a key mediator in hypothalamic-pituitary-adrenal signaling [13].

Multiple membrane trafficking proteins have been described as playing distinct roles in the sorting and processing of peptide hormones, however it remains unknown if these proteins affect the sorting of POMC to DCSGs. A critical membrane trafficking protein family is the heterotetrameric adaptor proteins (APs), AP-1 and AP-3, which mediate keys stages of DCSG formation and release [14]. Indeed, AP-1 is critical in selectively removing cargo from the maturing DCSGs, such as the vesicular associated membrane protein-4 (VAMP4) [15], calcium independent-mannose-6-phosphate receptor (CI-MPR) [16] and furin [17]. Furthermore, knockdown of the AP-3δ subunit in *C. elegans* results in defects in DCSG density, size, and number [18]. Interestingly, the regulation of AP-1 and AP-3 relies on their ability to interact with the multifunctional membrane trafficking regulator phosphofurin acidic cluster sorting protein 1 (PACS-1) [19]. Specifically, the PACS-1:AP-1 interaction is mediated through PACS-1 residues 174-182, and facilitates the removal of VAMP4 from immature secretory granules through the VAMP4 cytoplasmic tail motif (E_27_DDSDEEED) [15, 19]. Furthermore, sorting of the cellular endopeptidase furin away from maturing secretory vesicles is mediated through PACS-1 and AP- 1 [17]. Thus, through its ability to regulate AP-mediated trafficking, PACS-1 regulates multiple membrane trafficking steps within the cell, making it a hub for membrane trafficking, including cargo in the regulated secretory pathway.

Herein, we provide evidenced that identifies PACS-1 and AP-1 as key mediators in the sorting of POMC to the regulated secretory pathway. We observed that mutation of the PACS-1: AP binding site decreases the ability of PACS-1 to co-localize with ACTH-positive DCSGs. Furthermore, knockdown of PACS-1 and AP-1 decreases DCSG formation, decreases intracellular ACTH storage, and increases extracellular ACTH levels in the absence of stimuli. Importantly, upon knockdown of PACS-1 and AP-1, we observed an increase in ACTH localization to the constitutive secretory pathway.

## Materials And Methods

### Cell Culture

AtT-20 cells (ATCC; catalog #CCL-89) were cultured in complete Dulbecco’s Modified Eagle’s Medium (DMEM; Hyclone, Logan, UT) supplemented with 10% fetal bovine serum (Wisent, Montreal, Canada), 10μM L-Glutamine (HyClone), and 100μM penicillin and streptomycin (Hyclone). Cells were grown at 37 °C in the presence of 5% CO_2_ and sub-cultured in accordance with supplier’s recommendations.

### Plasmids and siRNA

Plasmids encoding VAMP2-GFP-C3 and GFP-tagged transferrin receptor were obtained from Thierry Galli (French Institute for Health and Medical Research; Addgene #42308) [20], and from Dr. Ron Flanagan (Western University). PACS-1 cDNA was provided by Dr. Gary Thomas (University of Pittsburgh), and sub-cloned into the pEGFP-N1 vector to yield PACS-1-GFP. Additionally, ADMUT-GFP was generated by PCR overlap mutagenesis. For knockdown experiments, siRNA against PACS-1 (CCUUAGCUGUGGGACUCAUTT), AP-1 μ-1 (GUAUCGGAAGAAUGAAGUATT), AP-3 δ-1 (GCGAUGAACUGCUCACCAATT) and a scrambled control were obtained from Life Technologies (Carlsbad, CA).

### Transfections

To quantify intracellular and extracellular levels of ACTH, 3 × 10^5^ AtT-20 cells were seeded into 12 well dishes and 24 hours later, cells were transfected with 2.5μM siRNA using PepMute^TM^ (FroggaBio, North York, Canada). Forty-eight hours post-transfection cells lysates and supernatants were collected for Western blot and MAGPIX analysis, respectively. For microscopy, 3 × 10^5^ AtT-20 cells were seeded on coverslips and transfected with 2.5μM siRNA and 1μg of the appropriate plasmid DNA using PepMute^TM^ siRNA or PolyJet Transfection Reagent (FroggaBio). Forty-eight hours post transfection, cells were fixed and prepared for imaging.

### qRT-PCR

AtT-20 cells were transfected with siRNA, as described above. Cells were collected 48 hours post transfection, and RNA was extracted using Ambion RNA extraction kit (Life Technologies). First strand cDNA synthesis was performed using the Superscript III RT cDNA synthesis kit (Life Technologies). Subsequently, cDNA was subjected to quantitative PCR using the SensiFast Probe Hi-Rox Kit (Bioline, Taunton, MA) and the Applied Biosystems QuantStudio 5 Real-time PCR system (Thermo Fisher Scientific, Waltham, Massachusetts, USA). Using *actin* as the reference gene, fold change in expression of the target gene was calculated using the delta CT (actin/target), and compared to that of scrambled control siRNA. Fold change expression = 1 - 2^-((CTactin_target_ – CTsiRNA_target_) - (CTactin_control_ – CTsiRNA_control_)).

### Western Blots

To assess intracellular protein levels in siRNA transfected cells, forty-eight hours posttransfection cells were lysed on ice with lysis buffer (0.5M HEPES, 1.25M NaCl, 1M MgCl_2_, 0. 25M EDTA, 0.1% Triton X-100, 1X complete protease inhibitor tablets (Roche, Indianapolis, IN)). and centrifuged at 4°C for 20 minutes Supernatants were collected and boiled for 10 minutes after addition of 5X SDS-PAGE sample buffer (5X Sample Buffer; 0.312M Tris pH 6.8, 25% 2-Mercaptoethanol, 50% glycerol and 10% SDS). Samples were run on a 14% SDS-PAGE gel and transferred onto a nitrocellulose membrane. Membranes were blocked in 5% skim milk powder in TBST for 1 hour, and incubated overnight at 4°C with antibodies specific for the ACTH peptide residues 1-18 (1:1000; provided by Dr. Iris Lindberg, University of Maryland) or actin (1:3000; Thermo Scientific). Membranes were washed 3X in TBST, incubated in species-specific HRP-conjugated secondary antibodies for 2 hours (1:2000; Thermo Scientific), and washed 3X in TBST. All blots were imaged using the C-Digit chemiluminescence Western blot scanner (LI-COR Biosciences, Lincoln, NE) with Crescendo ECL substrate (Millipore Inc.; Billerica, MA). Band intensities corresponding to peptides of 29 kDa, 26 kDa, and 9 kDa were quantified using the ImageQuant Studio software (LI-COR) and normalized to the actin loading control.

### Extracellular ACTH Quantification

To quantify basal extracellular levels of ACTH, cells were transfected with siRNA, as above. Forty-eight hours post transfection, cell culture media was removed and 250μl of fresh complete DMEM added. Cells were then incubated at 37°C for 2 hours. Subsequently, cell supernatants were collected and centrifuged at 10 000 × *g* to remove particulates and supernatants were subjected to MAGPIX analysis using the ACTH MILLIPLEX MAP mouse pituitary magnetic bead panel (Millipore, Etobicoke, Ontario).

### Immunofluorescence

#### Widefield imaging microscopy

Forty-eight hours post transfection, cells were washed 3X in PBS, fixed in 4% paraformaldehyde for 15 minutes, and subsequently washed 3X with PBS. Cells were blocked for 1 hour in blocking buffer (5% bovine serum albumin and 0.1% Triton X-100 in PBS). Cells were then incubated with rabbit anti-ACTH (1:200;) and/or mouse anti-LAMP1 (1:100; clone 1D4B; Developmental Studies Hybridoma Bank; Iowa, USA) for 2 hours. Cells were washed 2X and incubated with the goat-anti-mouse Cy3, or goat-anti-Rabbit Alexafluor 647 secondary antibody (1:800; Jackson ImmunoResearch) for 1.5 hours. Cells were then mounted on Fluoromount-G mounting media containing DAPI nuclear stain. Cells were viewed on a Leica DMI6000B (Leica, Wetzlar, Germany), and images were deconvolved using the advanced fluorescence application on the Leica Application Suite software. Co-localization analysis was performed using the Pearson’s correlation coefficient from the ImageJ plugin JACoP, as described previously [21].

#### Ground-state depletion imaging

Super-resolution imaging was performed as described previously [21]. Briefly, cells were mounted, and imaged in a depression slide containing 100mM cysteamine (Sigma Aldrich). All images were acquired using a Leica SR GSD microscope with a 100X/1.43 NA objective lens containing a 1.6X magnifier. GFP and AlexaFluor 647 fluorophores were excited using 488 nm and 647 nm laser lines respectively. Resulting images were quantified using the spatial association analysis MIiSR software, as previously described [21] (Available for download: http://phagocytes.ca/miisr/).

### Statistics

All statistics were conducted using the unpaired t-test or the one-way ANOVA with multiple comparisons where indicated using Graph Pad Prism (Graph Pad Software Inc., La Jolla, CA).

## RESULTS AND DISCUSSION

### PACS-1 and AP-1 promote intracellular storage of POMC

PACS-1 mediates trafficking of multiple proteins to and from the trans-Golgi network (TGN) through its ability to recruit AP-1 and AP-3 [19]. Thus, we sought to test for a role of PACS-1, AP-1, and AP-3 in the sorting of POMC to DCSGs by performing knockdown studies in mouse pituitary AtT-20 cells, which endogenously expresses POMC [6]. Transfection of siRNAs targeting the *PACS-1*, *AP-1*, or *AP-3* genes resulted in approximately 50% knockdown relative to non-specific siRNA, as determined by qRT-PCR (Fig. 1A). Subsequent to siRNA transfection, intracellular levels of POMC and its intermediates were assessed using Western blotting (Fig. 1B). Total intracellular POMC cleavage products were detected by Western blot using an antibody which detects residues 1-18 of ACTH, and were quantified using densitometry [22] (Fig. 1C). Strikingly, upon knockdown of PACS-1 or AP-1, we observed a 50% reduction in intracellular POMC-derived peptides compared to non-specific siRNA. However, knockdown of AP-3 did not significantly alter the intracellular storage levels of POMC (Fig. 1C). Interestingly, the processing of POMC was not affected when PACS-1 or Adaptor Protein levels were reduced (Fig. 1). This is consistent with the notion that PACS-1 may mediate sorting of POMC-derived peptides following its cleavage by a PC enzyme. Alternatively, upon depletion of PACS-1 and AP-1; PCs and POMC may mis-localize to the same compartment, wherein POMC can still be cleaved into active components, yet is secreted in an unregulated fashion.

**Figure 1:**
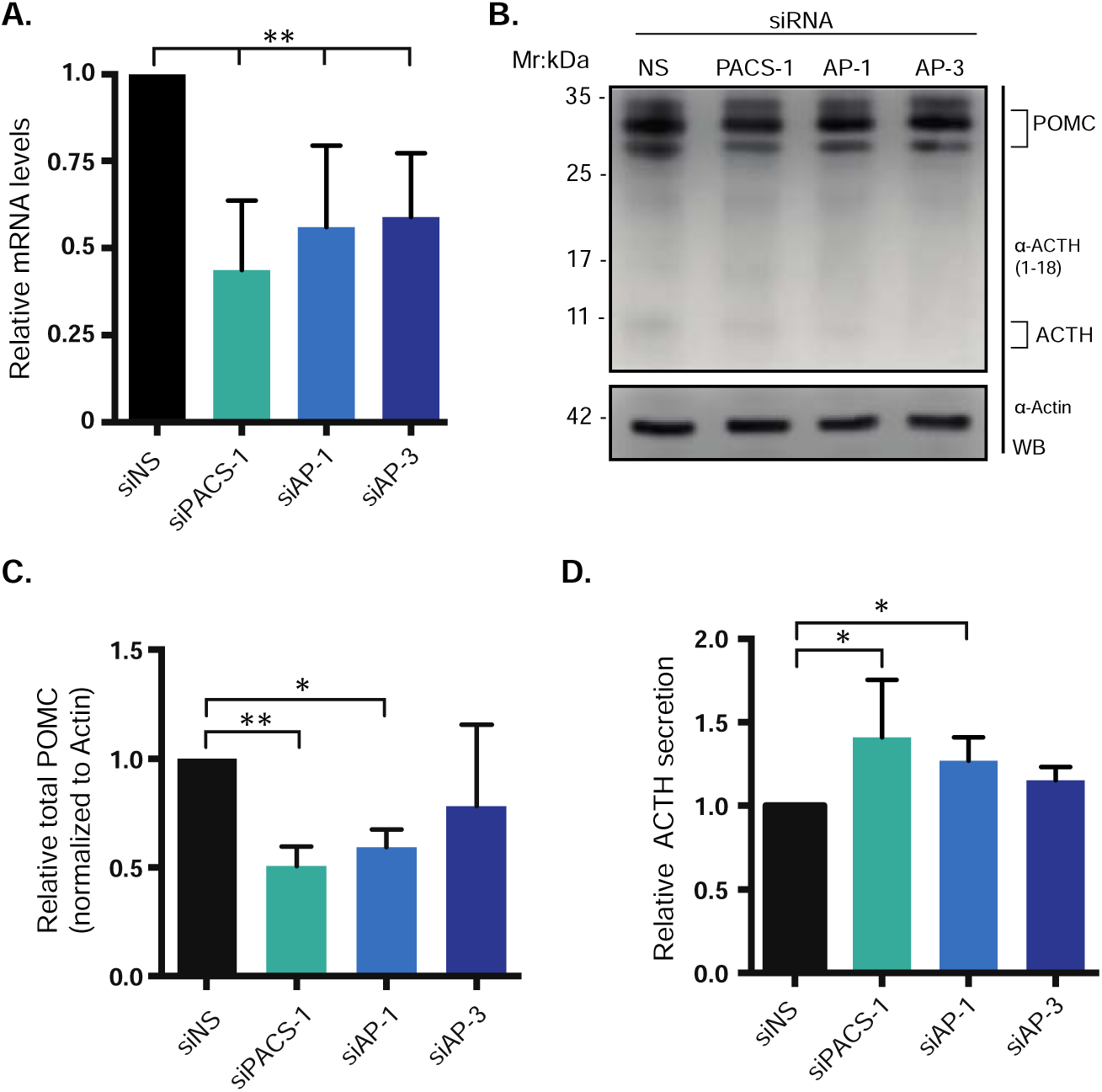
Intracellular storage of POMC is regulated by PACS-1 and AP-1. AtT-20 cells were transfected with siRNA targeting *PACS-1*, *AP-1*, *AP-3* or scrambled control. Forty-eight hours post transfection, cell-associated and cell culture supernatant-associated POMC was quantified. (A) Knockdown efficiency of siRNA transfections was determined by reverse transcription of cellular RNA to cDNA followed by quantification of siRNA target cDNA via qRT-PCR. Percent knockdown was calculated by comparing relative cDNA levels within cells transfected with targeting siRNA versus the scrambled control (n=6). (B) AtT-20 cell lysates were subjected to Western blot 48 hours post siRNA transfection with antibodies targeting ACTH 1-18, and the loading control Actin. (C) Quantification of mean (+/- standard error) intracellular POMC protein levels was completed byWestern blot. Levels of POMC were calculated relative to Actin and subsequently normalized to that of POMC in the scrambled control siRNA treated cells. Shown is the quantification of 5 independent experiments (n=5). (D) Supernatants of siRNA transfected cells were subjected to quantification of secreted extracellular ACTH using a MAGPIX assay. Mean relative extracellular ACTH (+/- standard error), compared to scrambled control siRNA, was calculated from 3 independent experiments (n=3). (* Indicates p-value <0.05, ** p-value <0.01; WB: Western blot).

We next sought to determine if knockdown of PACS-1, AP-1 or AP-3 resulted in the increased secretion of the POMC-derived peptide ACTH, in the absence of external stimulation. We observed 1.5 fold and 1.25 fold increases in ACTH in the cell culture supernatant upon knockdown of PACS-1 and AP-1, respectively, compared to the non-specific control siRNA by MAGPIX analysis (Fig. 1D). In contrast, increased levels of extracellular ACTH were not detected in cell culture supernatants from AP-3 knockdown cells (Fig. 1D). These findings indicate that upon knockdown of PACS-1 or AP-1, unregulated secretion of ACTH occurs, suggesting that sorting of POMC to the regulated secretory pathway is dependent on the membrane trafficking proteins PACS-1 and AP-1, but not necessarily AP-3.

To further assess the role of PACS-1 and its binding partners AP-1 and AP-3 in the sorting of POMC-derived peptides to DCSGs, we visualized the localization of ACTH in AtT-20 cells using widefield microscopy (Fig. 2A). Accordingly, we co-transfected siRNA against PACS-1, AP-1, AP-3 or a non-specific sequence with a plasmid expressing a fluorescently labeled marker of the unregulated secretory pathway, the transferrin receptor (TfnR) [23], and examined the cells via microscopy (Fig. 2A). We subsequently compared the co-localization of POMC/ACTH with TfnR-GFP [23] in the presence or absence of siRNA targeting the various trafficking proteins. Our results suggest that in PACS-1 knockdown cells, POMC/ACTH is shunted to the unregulated secretory pathway, as we observed an increased co-localization of POMC/ACTH with TfnR-GFP (siPACS-1; Fig. 2B; Pearson’s Coefficient = 0.49), compared to the scrambled siRNA control (siNS; Fig. 2B; Pearson’s Coefficient = 0.32). Similarly, knockdown of AP-1 or AP-3 resulted in a significant increase in the co-localization of ACTH with TfnR-GFP (Pearson’s Coefficient = 0.46, 0.49, respectively). Taken together, these results support the hypothesis that PACS-1 and APs play important roles in the trafficking of POMC/ACTH to the regulated secretory pathway.

**Figure 2:**
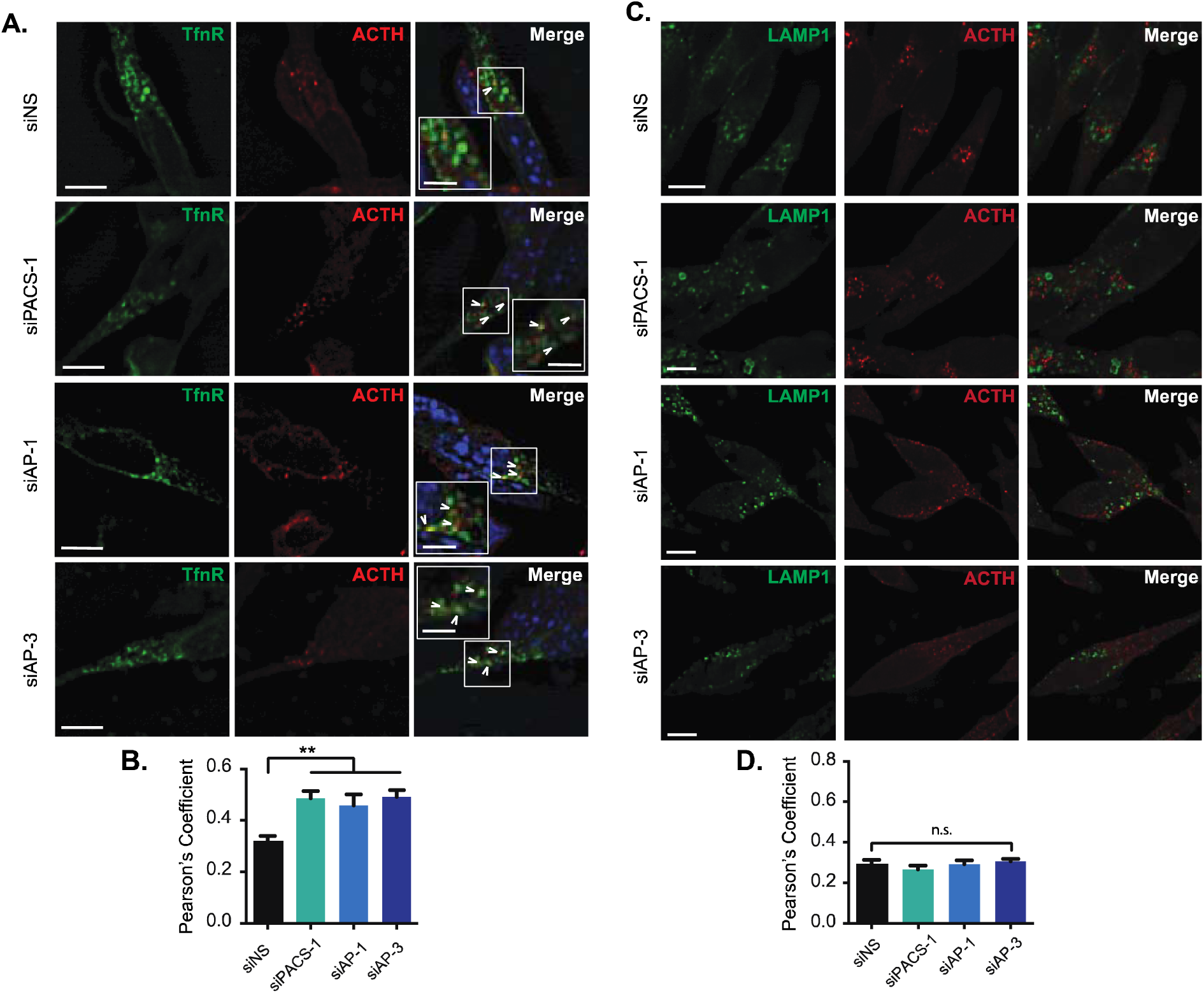
PACS-1, AP-1 and AP-3 are required for the sorting of ACTH to the regulated secretory pathway. AtT-20 cells were co-transfected with siRNA targeting *PACS-1*, *AP-1*, *AP-3* or a scrambled control and GFP-tagged transferrin receptor (TfnR-GFP). Forty-eight hours post transfection cells were immunostained for ACTH 1-18 and imaged via widefield microscopy. (A) Representative images of cells co-transfected with siRNA targeting *PACS-1* (siPACS-1), *AP-1* (siAP-1), *AP-3* (siAP-3) or a scrambled control (siNS), and TfnR-GFP. Shown are TfnR-GFP (Green), ACTH (Red) and DAPI (Blue). Scale bar represent 10μm, inset scale bars represent 5μm. (B) The co-localization between TfnR-GFP and ACTH was determined by calculating the mean (+/- standard error) Pearson’s correlation coefficient in at least 30 cells from 3 independent experiments. (** Indicates p-value > 0.01). (C) Representative images of cells transfected with siRNA targeting *PACS-1* (siPACS-1), *AP-1* (siAP-1), *AP-3* (siAP-3) or a scrambled control (siNS). Cells are stained for LAMP1 (Green) and ACTH (Red). Scale bar represent 10μm. (D) Co-localization between LAMP1 and ACTH staining was scored by calculating the mean (+/- standard error) Pearson’s correlation coefficient by quantification of at least 40 cells in 3 independent experiments. (n.s. represents not significant, LAMP1; Lysosomal associated membrane protein - 1).

### ACTH is not directed to a lysosomal degradation pathway

We next tested if the reduced intracellular storage of POMC-derived peptides upon PACS-1 and AP-1 silencing could be attributed to the trafficking of POMC molecules to a degradative lysosomal compartment. To assess this, we analyzed ACTH co-localization with the lysosomal marker LAMP-1 in the presence of PACS-1, AP-1, AP-3 or control siRNA (Fig. 2C). Co- localization analysis between LAMP-1 and ACTH revealed that decreased expression of PACS-1, AP-1 or AP-3 does not result in the trafficking of ACTH to a lysosomal compartment, compared to control siRNA (Fig. 2D). Taken together, these results suggest that the observed decrease in intracellular storage of POMC-derived peptides is not driven through the degradation of ACTH.

### PACS-1: Adaptor Protein interactions modulate PACS-1:ACTH co-localization

Both PACS-1 and AP-1 act in concert to recognize cargo and modulate their localization within cells. For example, expression of a dominant-negative form of PACS-1, which is defective in AP-1/3 binding (termed ADMUT; **Ad**aptor Protein binding **Mut**ant), alters AP-1 localization, preventing furin trafficking from endosomes to the TGN [19]. Therefore, we sought to determine if PACS-1 co-localized with POMC-positive vesicles in an AP-dependent manner. To test this, we examined co-localization between PACS-1/ADMUT and ACTH through super-resolution ground-state depletion microscopy (GSDM) using nearest neighbor spatial association analysis [24, 25] (SAA; Fig. 3A, B and C). GSDM allows for a 10-fold increase in resolution compared to conventional widefield or confocal microscopy [25]. Interestingly, we observed that over 60% of ACTH co-localized with PACS-1-GFP (Fig. 3A and D). Furthermore, SAA revealed that this co-localization occurred at a much greater rate than expected of randomly positioned molecules, suggesting that PACS-1 is within close proximity to ACTH vesicles, whereby it may be exerting its sorting function (Fig. 3C and D). Upon expression of ADMUT-GFP we observed a 30% reduction in co-localization of ACTH and ADMUT-GFP compared to wildtype PACS-1-GFP, as determined via super-resolution nearest-neighbor analysis (Fig. 3B and C). These results are akin to previous findings which demonstrate PACS-1 as an important regulator furin trafficking to and from immature secretory granules in an AP-1 dependent manner [26].

**Figure 3:**
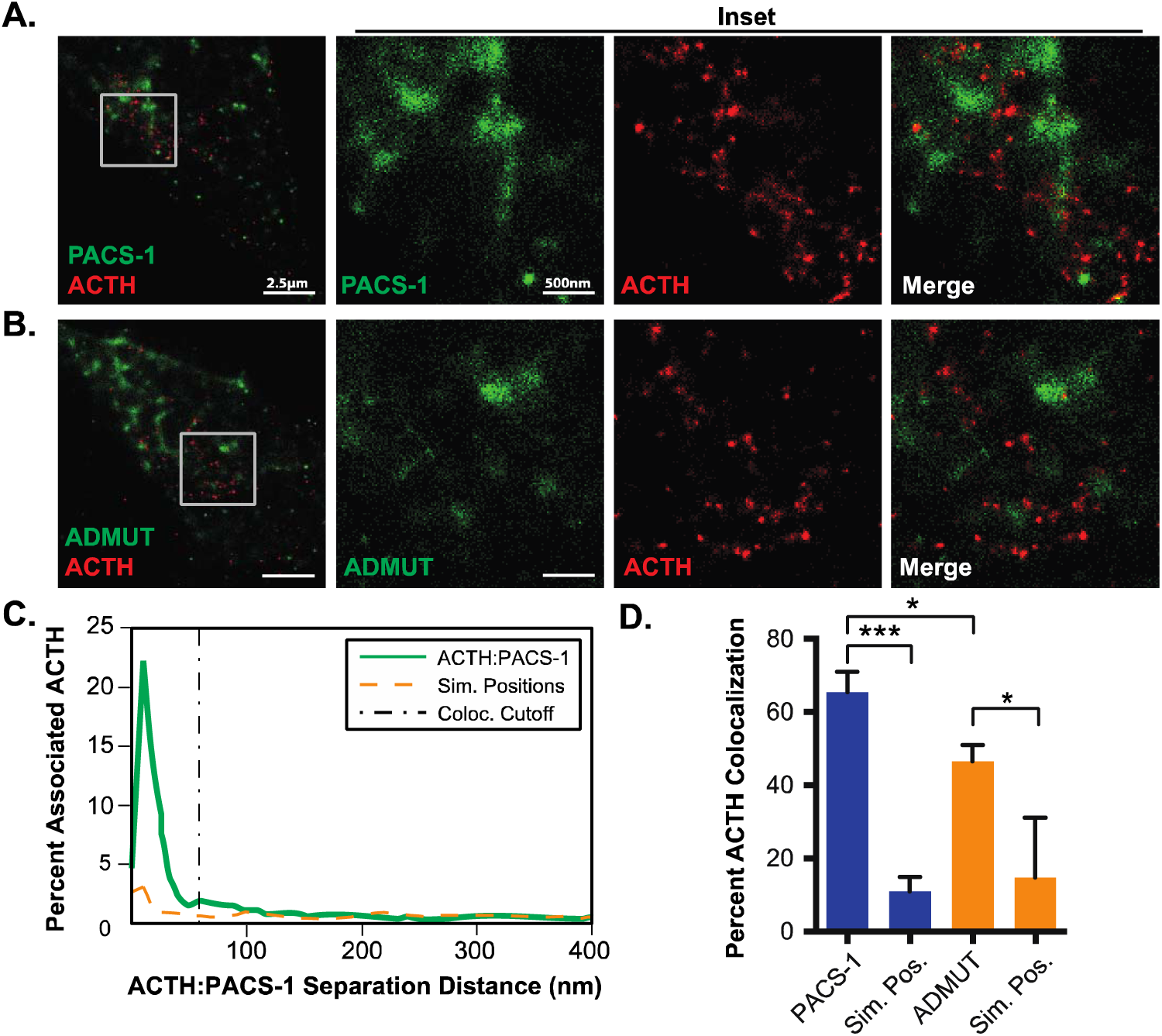
PACS-1 co-localizes to POMC^+^ vesicles in AtT-20 cells. AtT-20 cells were transfected with PACS-1-GFP or ADMUT-GFP and forty-eight hours post transfection, cells were fixed, immunostained for ACTH, and imaged using ground-state depletion super-resolution microcopy. (A+B) Representative images of AtT-20 cells transfected with PACS-1-GFP (A; Green) or ADMUT-GFP (B; Green) and immunostained for ACTH (Red) (scale bars represent 2.5μm, inset scale bars represent 500nm). (C) A representative histogram mapping the intermolecular distances between ACTH and PACS-1 (Green) or simulated random position (Orange). Co-localization cut-off is illustrated (black dashed line). (D) A graphical representation of the mean (+/- standard error) percent of ACTH co-localized with PACS-1 or ADMUT-GFP, or ACTH co-localized to randomly simulated positions. Error bars were calculated by quantification of 6 cells in 2 independent experiments. (* p-value < 0.05; *** p-value < 0.001)

### PACS-1 and AP-1 facilitate the trafficking of ACTH to DCSGs

We next sought to define if PACS-1 and APs specifically mediate the targeting of ACTH to DCSGs. To ascertain this, we used the vesicular associated membrane protein 2 (VAMP2) as a marker for DCSGs [27]. Due to the observed increase in constitutive secretion of ACTH upon knockdown of PACS-1 and AP-1 (Fig. 1D), we investigated the ability of ACTH to be targeted to VAMP2 positive DCSGs in the presence or absence of siRNA targeting PACS-1, AP-1 or AP- 3 (Fig. 4A). Strikingly, upon knockdown of PACS-1, we observed a significant reduction in the ability of ACTH (red) to be localized to VAMP2-positive compartments (green) compared to the non-specific siRNA control (Fig. 4A and B), highlighting the role of PACS-1 in sorting ACTH to mature granules. Furthermore, knockdown of AP-1 but not AP-3 resulted in an even greater reduction of ACTH localization to VAMP2-positive compartments (Fig. 4A and B). Taken together, the results demonstrate the importance of PACS-1 and AP-1 in sorting ACTH containing vesicles to mature secretory granules.

**Figure 4:**
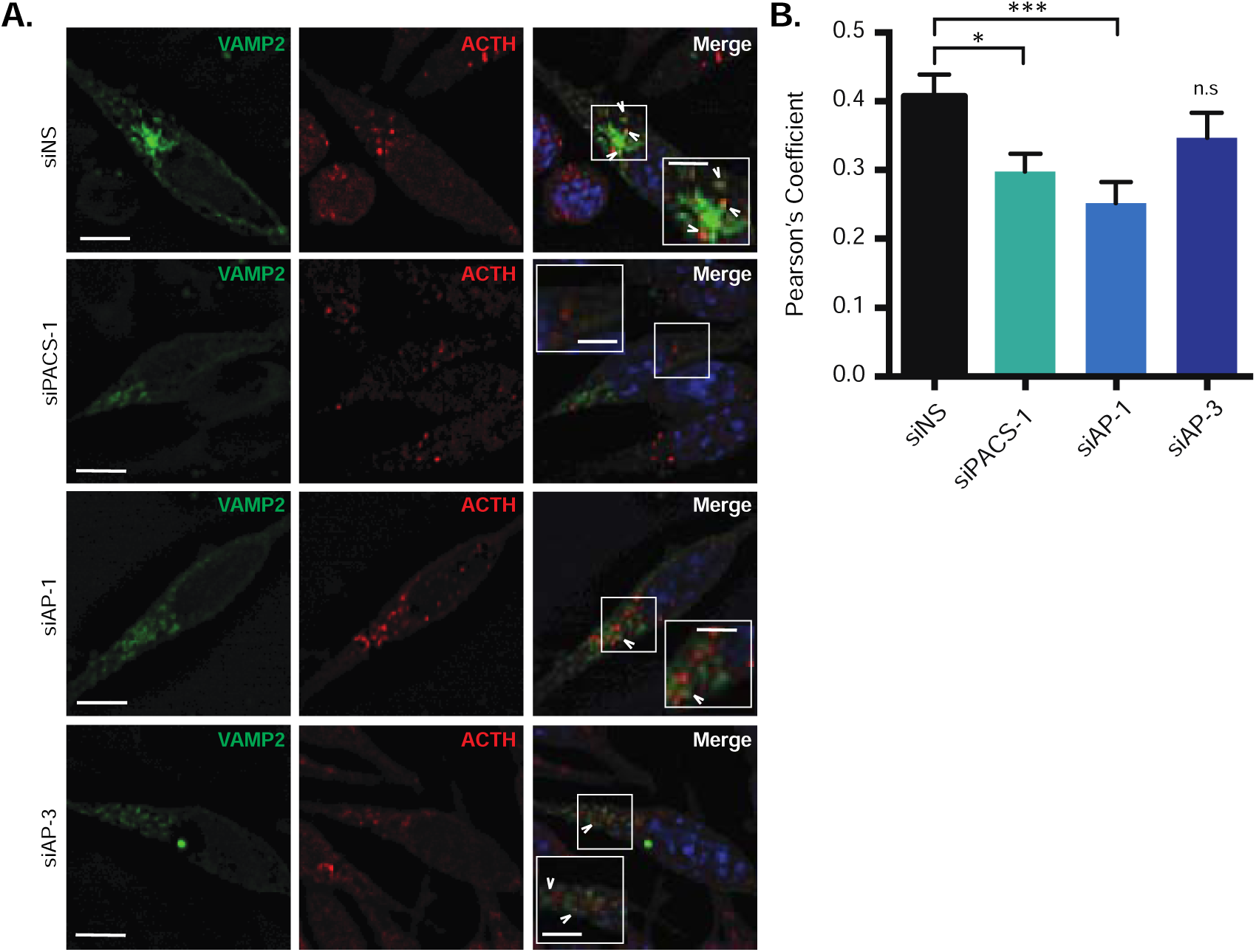
PACS-1 and AP-1 facilitate the sorting of POMC to mature secretory granules. AtT-20 cells were co-transfected with siRNA against *PACS-1*, *AP-1*, *AP-3* or scrambled control and a plasmid encoding GFP-tagged Vesicular Associated Membrane Protein – 2 (VAMP2- GFP). Forty-eight hours post transfection cells were fixed and immunostained for ACTH. (A) Representative images illustrating VAMP2-GFP (Green), ACTH (Red) and DAPI (Blue) in cells transfected with siRNA targeting *PACS-1* (siPACS-1), *AP-1* (siAP-1), *AP-3* (siAP-3) or a scrambled control (siNS). Scale bar represent 10μm, inset scale bars represent 5μm. (B) The mean (+/- standard error) co-localization between ACTH and VAMP2-GFP was scored by calculating the Pearson’s correlation coefficient using the JACoP plugin on ImageJ from the quantification of at least 25 cells from 3 independent experiments. (* indicates p-value < 0.05; *** p-value < 0.001; n.s. represents not significant).

Interestingly, knockdown of AP-3 has been associated with the routing of the DCSG-resident protein, secretogranin II, to the unregulated secretory pathway [18]. In agreement with these findings, we observed that knockdown of AP-3 resulted in an increase in ACTH localization to the constitutive pathway (Fig. 2A). However, we did not observe a corresponding decrease in sorting of ACTH into VAMP2 positive granules (Fig. 4B), nor did we observe an increase in extracellular ACTH secretion upon AP-3 knockdown (Fig. 1D). It is possible that AP-3 knockdown results in the compensation by other APs, or that ACTH does not follow the AP-3 dependent pathway previously described, as not all DCSGs contain the same cargo [28].

Overall, we provide evidence supporting the key role of PACS-1 and AP-1 in controlling peptide hormone storage within AtT-20 cells. Controlling the release of peptide hormones is an important physiological process, which requires multiple proteins. Studying how these proteins work together is fundamental to understanding how different pathways can be altered in homeostasis and disease.

## ACKNOWLEDGEMENTS

We acknowledge Drs. Gary Thomas, Ron Flannagan, Iris Lindberg and Thierry Galli for reagents. This work was supported by a NSERC Discovery Grant to JDD (grant #435677). Work in the BH lab was supported by (NSERC grant #418194). BSD is supported by a CIHR Doctoral Fellowship, ENP is supported by an NSERC Doctoral Fellowship. CE was supported by an NSERC USRA.

## AUTHOR CONTRIBUTIONS

BSD, CE and, LVN preformed experiments. BSD, JDD, ENP conceived and designed experiments. BSD, CE, ENP, BH, RAJ and JDD analyzed data. All authors aided in the writing of the manuscript.

